# STAT3 expression is reduced in cardiac pericytes in HFpEF and its loss reduces cellular adhesion and induces pericyte senescence

**DOI:** 10.1101/2025.02.14.638233

**Authors:** Leah Rebecca Vanicek, Ariane Fischer, Mariano Ruz Jurado, Tara Procida-Kowalski, Jochen Wilhelm, Maximilian Merten, Felicitas Escher, Badder Kattih, Valentina Puntmann, David John, Eike Nagel, Stefanie Dimmeler, Guillermo Luxán

**Affiliations:** Institute of Cardiovascular Regeneration, Goethe University Frankfurt, Theodor Stern Kai 7, Frankfurt am Main, Germany; Cardiopulmonary Institute, Frankfurt am Main, Germany; DZHK, site Rhein/Main, Frankfurt am Main, Germany; Institute for Lung Health. Justus-Liebig-University Giessen, Aulweg 132, Giessen, Germany; Universities of Giessen and Marburg Lung Center (UGMLC), German Center for Lung Research (DZL), Justus-Liebig University Giessen, Giessen, Germany; Institute of Cardiac Diagnostics and Therapy, IKDT GmbH Berlin, Germany; Department of Cardiology, University Hospital Frankfurt, Goethe University Frankfurt. Frankfurt am Main, Germany; Institute for Experimental and Translational Cardiovascular Imaging, DZHK Centre for Cardiovascular Imaging, Goethe University Frankfurt, Germany

## Abstract

Heart failure with preserved ejection fraction (HFpEF) accounts for half of heart failure cases and is characterised by reduced pericyte coverage. While the contributions of other cardiac cell types to HFpEF are well-studied, the role of pericytes remains less understood. Using murine single-nucleus RNA sequencing to study cardiac pericytes in HFpEF, we identified reduced *STAT3* expression as a hallmark of HFpEF pericytes. Mechanistic studies *in vitro* revealed that *STAT3* deletion induces cellular senescence and impairs pericyte adhesion, recapitulating HFpEF-like characteristics. These findings suggest that *STAT3* is crucial for maintaining pericyte homeostasis and highlight its reduction as a potential driver of pericyte loss, a defining feature of HFpEF.

## Introduction

Heart failure with preserved ejection fraction (HFpEF), referred to as diastolic heart failure, accounts for half of heart failure patients [1,2]. Ageing, a primary risk factor for cardiovascular disease, is strongly linked to HFpEF development [3]. HFpEF is characterized by stiffening of the left ventricle and an increase in left ventricular wall thickness, leading to increased end-diastolic pressure, hinders adequate left ventricular filling [1,4] and is presented with a left ventricular ejection fraction higher than 50% [5–7]. Despite being associated with high morbidity and mortality, only recently, however, certain treatments, such as SGLT2 inhibitors, have shown efficacy in reducing hospitalisation rates and cardiovascular death risk [8,9].

Microvascular dysfunction is a key cellular mechanism underlying HFpEF [10,11]. Studies using a rat HFpEF model suggest that pericyte loss initiates microvascular dysfunction, contributing to diastolic dysfunction [12]. Pericytes are capillary-associated mesenchymal cells that play an important role during angiogenesis, vascular permeability control, extracellular matrix deposition, and blood-brain-barrier maintenance [13,14]. In the heart, pericyte dysfunction can lead to interstitial fibrosis, atherothrombosis, or calcification vasculopathy [15], furthermore, age-related pericyte dysfunction has been shown to induce myocardial fibrosis, a hallmark of HFpEF [16]. While the roles of cardiomyocytes, endothelial cells, and fibroblasts in HFpEF-related remodelling have been well-studied, the molecular and cellular changes in pericytes during HFpEF progression remain largely unknown.

This study identifies the loss of the transcription factor *STAT3* (Signal Transducer and Activator of Transcription 3) in cardiac pericytes, observed in both human HFpEF patients and a mouse HFpEF model, as a marker of pericyte dysfunction. Furthermore, our findings demonstrate that STAT3 is essential for maintaining pericyte homeostasis, as its loss impairs cellular adhesion and induces cellular senescence.

## Material and Methods

### Mice

Male and female C57Bl/6J mice were used in this study. Mice were purchased from Janvier (Le Genest Saint-Isle, France). To induce HFpEF, mice were exposed to a combination of high-fat diet (60 % kilocalories from fat (lard)) and N[w]-nitro-l-arginine methyl ester (L-NAME; 0.5 g/l in drinking water, Sigma-Aldrich) for 10 weeks as described in [17,18]. Animals were held at 23 °C ambient temperature and 60% humidity in 12 h/12 h light/dark cycle. Animal experiments were performed in accordance with the principles of laboratory animal care as well as German national laws, the Directive 2010/63/EU of the European Parliament, and were approved by the ethics committee of the regional council (Regierungspräsidium Darmstadt, Hessen, Germany) under the animal experiment license FU/1247. At the end of each experiment, animals were euthanised by cervical dislocation after isoflurane administration (2-2,5 Vol.-%).

### Human samples

Human cardiac biopsies were obtained from the study Decipher-HFpEF study (NCT03251183). Two healthy control biopsy were kindly donated by Prof. Dettmeyer from the University Gießen. This is a post-mortem sample with unrelated cause of death. The cardiac biopsies were examined to discard any pathological alterations. The cardiac biopsies were fixed and embedded in paraffin. Paraffin blocks were sectioned in 4 μm thick sections on a microtome (HM340, ThermoFisher), transferred to microscope slides and stored at room temperature.

### Immunofluorescence staining on paraffin sections

Paraffin slides were prewarmed at 60 °C for 30 min and deparaffinized with two consecutive 10 min washes with rotihistol. Next, slides were rehydrated with a series of ethanol washes in decreasing concentration, 100%, 95%, 80%, 70%, 50% ethanol and distilled water for 5 min each. After that, antigen retrieval was performed in 0.01 M citrate buffer (pH=6.0) for 90 sec in a pressure cooker and cooled down by rinsing with tab water for 2 to 3 min. Sections were washed three times for 5 min in PBS containing 0.1% Triton and blocked with blocking solution (3 % BSA, 20 mM MgCl_2_, 0.1% TritonX-100, 5% donkey serum, diluted in PBS) for 1 h at room temperature in a humid chamber. Samples were incubated with primary antibodies in blocking solution in a humid chamber over night at 4 °C. The next day, slides were washed three times for 5 min in PBS and incubated with secondary antibodies diluted in PBS containing 5% BSA for 1 h at room temperature in the dark in a humid chamber. Samples were washed twice for 5 min with PBS and once for 5 min in PBS containing 0.1% Triton. Finally, slides were mounted with Fluoromount-G (00-4958-02, ThermoFisher). After letting the mounting medium cure at room temperature, samples were stored at 4°C. Imaging was performed with a Leica Stellaris confocal microscope and quantification was done with Volocity software.

#### Primary antibodies and dyes

Rabbit anti-NG2 (1:50; PA592029, Invitrogen); goat anti-PDGFRβ (1:50; GT15065, Neuromics); Biotinylated Griffonia Simplicifolia Lectin I (GSL I) isolectin B4 (1:50; B-1205, Vector Laboratories); Biotinylated Ulex Europaeus Agglutinin I (UEA I) (1:50; B-1065-2, Vector Laboratories).

#### Secondary antibodies

Donkey anti-rabbit Alexa Fluor 488 (1:200; A-21206, Invitrogen); Donkey anti-goat Alexa Fluor 555 (1:200; A-21432, Invitrogen). 4’, 6-diamidino-2- phenylindole (DAPI) (1:1000; D9542, Sigma) was used to counterstain the nuclei.

### Single-nucleus-RNA-sequencing

The single-nucleus-RNA-sequencing data set generated in [18] was used in this study.

### Cell culture

Human placenta pericytes (hPL-PC, C-12980, PromoCell) were cultured in Pericyte Growth Medium 2 (C-28041, PromoCell) at 37 °C and 5% CO2. When cells reached around 80-90% confluency, they were split using Accutase (#A6964, Abcam). All experiments were performed with cells between passages 3 and 6.

### STAT3 knockdown

*STAT3* silencing was performed using siRNA (Sigma; sequences: GGAUAACGUCAUUAGCAG, UCUGCUAAUGACGUUAUCC. Concentration, 100µM). hPL-PC were seeded with a density of 77.000 cells/well in a 6-well-plate 24 h before transfection. STAT3 siRNA and control siRNA were used at a final concentration of 50 nM. For each well an independent solution, containing siRNA diluted in 200 µl OptiMEM medium (51985034, Gibco) was prepared. Additionally, 5 µl Lipofectamine RNAiMAX transfection reagent (13778150, ThermoFisher) was diluted in 195 µl OptiMEM. The siRNA diluted in OptiMEM was added to the Lipofectamine mix and incubated for 15 min at room temperature. Cells were washed with 1 ml OptiMEM medium per well. Then, 1.6 ml OptiMEM medium was added per well and the transfection mix (400 µl) was added dropwise. After 4 h incubation the medium containing the transfection reagents was removed, and cells were further cultured in Pericyte Growth Medium 2 (C-28041, PromoCell) as indicated above.

### Migration assay

Cells were seeded in 2-well inserts (80209, IBIDI). The 2-well inserts were placed in each well of a 12-well-plate in technical triplicates per condition. Transfected cells were washed with PBS and detached using 500 µl Accutase per well (of a 6-well plate) for 4 min at 37 °C. Cell number was determined, and cells were reseeded at a density of 25.000 cells/insert side in a total volume of 70 µl. 1 ml Pericyte Growth Medium was added around the Insert. Cells were incubated for 24 h and after that the insert was removed. The medium was then removed, and 1 ml fresh medium was added per well. Images of the cell-free gap were taken at time point 0 (directly after removing the insert) 8h and 16h after removing the insert. The cell free area was analyzed with ImageJ and the migration capacity was calculated as percentage of closed area from time point 0.

### Adhesion assay

Adhesion was investigated by the capacity of pericytes to attach to a gelatin-matrix. To do so, three wells per sample of a 96-well plate were coated with 200 µl gelatin (1:10 in PBS) and incubated for 1 h at 37 °C. After that, wells were washed twice with 100 µl supplement free medium + 0.05% BSA and 50 µl supplement free medium + 0.05% BSA was added per well. Cells were stained with 0.26 µl BCECF-AM (1 mg/ml; #B1150, ThermoFisher) in 1 ml supplement free pericyte medium for 30 min at 37 °C. Cells were detached using Accutase and centrifuged at 700 g for 5 min. Cell number was adjusted to 1x10^6 cells/ml and 50 µl of cell solution was seeded in triplicates in gelatin coated wells. Cells were incubated for 1 h at 37°C to let the cells adhere to the gelatin surface. Next, fluorescence intensity was measured at 535nm with Multi-Detection Microplate Reader Synergy HT (BioTek). Cells were washed eight times with 100 µl supplement free pericyte medium + 0.05% BSA and fluorescence intensity was measured after each washing step. Adhesion was calculated as measurement 2 (after the first washing step) divided by measurement 1 (before the first washing step).

### Immunocytochemistry

Cells were reseeded in a density of 12.500 cells/well into µ-Slide 4 Well slide (80426, Ibidi) coated with gelatin and incubated overnight at 37°C. On the next day, cells were washed three times for 5 min with PBS. Cells were fixed with 4% PFA in PBS for 15 min and washed three times with PBS for 5 min. Cells were permeabilized with 0.1% Triton/PBS for 10 min. Next, blocking of non-specific antibody interactions was performed using PBS containing 5% donkey serum for 60 min at room temperature and incubated with primary antibodies diluted in blocking solution over night at 4 °C. On the next day, cells were washed three times for 5 min with PBS and incubated with secondary antibodies diluted in PBS for 60 min at room temperature. Cells were washed three times with PBS for 5 min and mounted with Fluromount-G (00-4958-02, ThermoFisher). After letting the mounting medium cure at room temperature, samples were stored at 4 °C. Imaging was performed with a confocal microscope using a 63x objective. Quantification was done with Volocity software.

#### Primary antibodies and dyes

Mouse Anti-Actin α Smooth Muscle - Cy3 (1:300; C6198, Sigma); Mouse anti-Vinculin (1:150; V9131, Millipore); Phalloidin Alexa Fluor 488 (A12379, Invitrogen)

#### Secondary antibodies

Donkey anti-mouse Alexa Fluor 647 (1:200; A-31571, Invitrogen). 4’, 6- diamidino-2-phenylindole (DAPI) (1:1000; D9542, Sigma) was used to counterstain the nuclei.

### CCK-8 assay

Cell Counting Kit-8 (CCK-8; #CK04-11, Dojindo) colorimetric assay was used to measure cell viability of pericytes. For this test, cells were reseeded at a density of 10.000 cells/well into a 96-well-plate. Each condition was pipetted in technical triplicates. One day after reseeding, the medium was changed to 90 µl OptiMEM medium (51985034, Gibco) per well. As a blank control 90 µl of OptiMEM medium were pipetted in triplicates in empty wells. Then, 10 µl CCK-8 solution was added per well and the plates were incubated for 2 h at 37 °C in the dark. After incubation, the absorbance was measured at a wavelength of 450 nm using Multi-Detection Microplate Reader Synergy HT (BioTek). The final absorbance was calculated as the difference between the measured absorbance and the mean of the blank.

### Acidic β-Galactosidase staining

Cellular senescence was studied using the Senescence β-Galactosidase cell staining kit (#9860, cell signaling). To do so, we seeded cells on 8-well µ-slides (80826, Ibidi) with at a density of 50.000 cells/well. Then, cells were washed twice with PBS for 5 min and incubated with 1x fixative solution for 30 min at room temperature. Cells were washed three times in PBS for 5 min. 10x staining solution was heated to 37°C with agitation and diluted to a 1x solution with distilled water. β-Galactosidase staining solution was prepared using 930 µl 1x staining solution, 10 µl 100x solution A, 10 µl 100x solution B and 50 µl 20mg/ml X-gal stock solution per sample. The pH value was adjusted to 6.0. Cells were incubated with β-Galactosidase staining solution over night at 37°C. On the next day, cells were washed two times with PBS for 5 min and after that, mounted with Pertex mounting medium (#41-4011-00, Medite). β-Galactosidase staining was imaged with Leica Eclipse TS100 microscope and analyzed with Volocity software.

### Telomere length

Genomic DNA was isolated using DNeasy blood and tissue kit (69504, Qiagen) according to manufacturer’s instructions. Telomere length was determined with qPCR using 5 µl SYBR Green Master Mix (4385617, Applied Biosystems), 1 µM primer, 0.5 µl H_2_O and 2.5 µl DNA per sample. The primers were designed to specifically amplify the telomere sequence (*TEL1*; forward: CGGTTTGTTTGGGTTTGGGTTTGGGTTTGGGTTTGGGTT, and reverse: GGCTTGCCTTACCCTTACCCTTACCCTTACCCTTACCCT) or a single-copy control gene (*36B4*; forward: CAGCAAGTGGGAAGGTGTAATCC, and reverse: CCCATTCTATCATCAACGGGTACAA). This allows to calculate the telomere length relative to the single copy control gene [19].

### BrdU assay

Cell cycle phases were analyzed using BrdU Flow kit (51-2354AK, BD Biosciences). hPL-PC were treated with 10 µM BrdU in culture medium for 4 h at 37°C. Cells were detached using 500 µl Accutase/well and centrifuged at 800g for 5 min. Cell pellet was resuspended in 100 µl Cytofix/Cytoperm buffer and incubated at room temperature for 15 min. Perm/Wash buffer was diluted 1:10 in distilled water. 1 ml Perm/Wash buffer was added to the samples. Samples were then resuspended, and centrifuged at 8000 rpm for 3 min. The cell pellet was resuspended in 100 µl Cytofix/Cytoperm buffer and incubated for 10 min on ice. Next, the cells were washed with 1 ml Wash buffer again and centrifuged at 8000 rpm for 3 min. The cell pellet was resuspended once more in 100 µl Cytofix/Cytoperm buffer and incubated on ice for 5 min. After that. The cells were washed with 1 ml Wash buffer and centrifuged at 8000 rpm for 3 min and then were incubated with 300 µg/ml DNase I for 1 h at 37°C, washed with 1 ml Perm/Wash buffer and centrifuged at 8000 rpm for 3 min. The cells were then stained with 2.5 µl mouse V450 anti-BrdU antibody (560810, BD) in 50 µl Perm/Wash buffer for 20 min at room temperature. Cells were washed with 1 ml Perm/Wash buffer and centrifuged at 8000 rpm for 3 min. Finally, 10 µl 7-AAD was added and incubated for 10 min at room temperature. After that 300 µl PBS were added, samples were transferred into FACS tubes and measured with BD Canto II FACS and Diva software. The results were analyzed with FlowJo software.

### Bulk RNA sequencing

Cells were transfected as indicated above and RNA was isolated using RNeasy Plus Mini Kit (74136, Qiagen) according to manufacturer‘s instructions. For whole-transcriptome analysis ribosomal RNA (rRNA) was removed from a total amount of 250 ng RNA per sample followed by cDNA sequencing library preparation utilizing the Illumina® Stranded Total RNA Prep, Ligation with Ribo-Zero Plus kit (Illumina) according to the manufacturer’s instructions. After library quality control by capillary electrophoresis (4200 TapeStation, Agilent), cDNA libraries were sequenced on the Illumina NovaSeq 6000 platform generating 50 bp paired-end reads.

### Real time quantitative polymerase chain reaction (RT-qPCR)

RNA was isolated using RNeasy Plus Mini Kit (74136, Qiagen) according to manufacturer‘s instructions. 500 ng of RNA were then reverse transcribed into cDNA using M-MLV reverse transcriptase (Thermo-Fisher Scientific). Quantitative PCR reactions were performed using the StepOnePlus real-time PCR cycler (Thermo-Fisher Scientific) and relative gene expression was calculated by normalisation to the average of *RLP0* and *ACTB* gene expression (2^-ΔCt^).

#### Primers

*ACTB* (f: CATGTACGTTGCTATCCAGGC; r: CTCCTTAATGTCACGCACGAT); *RLP0* (f: TCGACAATGGCAGCATCTAC, r: ATCCGTCTCCACAGACAAGG); STAT3 (f: CAGCACCTTCAGGATG, r: GCTTGACTCTTGAGGGTTT); *COL1A1* (f:GTTCGTGACCGTGACCTCG, r: TCTTGTCCTTGGGGTTCTTGC); COL3A1 (f: TCCTGGGAGAAATGGTGACC; r: GCGAGTCCTCCTACTGCTAC); *CSPG4* (f: GAGCCCAGGCACGAAAAATG, r: GTATGTTTGGCCCCTCCGAA); *PDGFRB* (f: AGCACCTTCGTTCTGACCTG, r: TATTCTCCCGTGTCTAGCCCA); *DES (f: ACATTTCTGAAGCTGAGGAG, r: GCGTCGTTGTTCTTGTTG); CDKN1A (f: AGTCAGTTCCTTGTGGAGCC, r: CATTAGCGCATCACAGTCGC)*

### Graphical figures

Graphical figures were originally created with BioRender.com or adapted from BioRender.com templates with modifications to original content and/or design using (licence number: 28D5A348-0001).

### Statistics

Statistical analysis was performed using GraphPad Prism 9.2.0 software. The data was tested for normal distribution (Shapiro-Wilk test) and after that an unpaired, two-tailed Student’s t-test was performed to compare two normally distributed groups. For not normally distributed groups, Mann-Whitney test was performed. For comparison of more than two groups a one-way analysis of variances (ANOVA) was performed for normally distributed groups followed by a post hoc Tukey’s test for multiple comparison. Data are presented as mean + standard error of the mean (SEM). All calculations were performed in GraphPad Prism 9.3.0.

### Data availability

Bulk-RNA-sequencing data will be made available upon acceptance

## Results

Histological analysis of cardiac biopsies from HFpEF patients (baseline characteristics in **Table 1**) revealed microvascular dysfunction characterized by increased capillary perimeter (**Figure 1A**) and reduced pericyte coverage in the myocardium (**Figure 1B**). Similarly, decreased pericyte coverage was observed in a murine model of HFpEF [17] (**Figure 1C**). To study the transcriptional effects of HFpEF on cardiac pericytes, we analyzed single-nucleus RNA sequencing from murine HFpEF samples [18] (**Figure 1D**). This dataset showed a well-defined pericyte cluster, marked by expression of canonical pericyte markers, including *PDGFRB*, *RGS5*, and *ABCC9* (**Supplementary Figure 1A**). Differentially expressed gene (DEG) analysis between control and HFpEF pericytes identified 2078 DEGs (P<0.05; logFoldChange>0.025), including 381 upregulated and 1697 downregulated genes. Gene ontology (GO) analysis revealed that genes related to terms like mitochondrial ATP synthesis, translation, and focal adhesion, were downregulated in cardiac pericytes (**Figure 1E**), while genes related to response to TGF-β, muscle contraction, and actin cytoskeleton were upregulated (**Supplementary Figure 1B**). To identify genes potentially driving pericyte dysfunction in HFpEF, we focused on dysregulated transcription factors that could be responsible for pericyte biology in the heart. STAT3 expression is reduced in cardiac murine HFpEF pericytes (**Figure 1F**) and its expression is also reduced in a cohort of human HFpEF patients [20] (**Figure 1G**). While STAT3’s role has been studied in cardiomyocytes and endothelial cells [21,22], its role in cardiac pericytes, particularly in the context of HFpEF, remains unknown.

**Table 1.** Baseline characteristics of patient cohort.

|  | **Control** | **HFpEF patients** |
| --- | --- | --- |
| **n** | 4* | 7 |
| **Age (average years)** | 44 | 64 |
| **Sex (male/female)** | (3/1) | (1/6) |
| **NYHA class [I/II/III/IV]** | [1/1/0/0] | [0/3/4/0] |
| **Hypertension [n (%)]** | 1 (50) | 6 (85.7) |
| **Diabetes [n (%)]** | 1 (50) | 2 (28.6) |
| **Smoking [n (%)]** | 0 (0) | 1 (14.3) |
| **Left ventricular EF (%)** | 53.5 | 59.3 |
| **ACEi or ATRB (%)** | 1 (50) | 6 (85.7) |
| **Beta-blocker (%)** | 0 (0) | 1 (14.3) |
| **Diuretic (%)** | 0 (0) | 2 (28.6) |
| **Aldosterone antagonist** | 0 (0) | 0 (0) |
| **Statin** | 1 (50) | 4 (57.1) |

\*besides age and sex, this information was known only for 2 of the patients
EF – ejection fraction
ACEi – angiotensin-converting enzyme inhibitor
ATRb – angiotensin II receptor blocker

**Figure 1.**
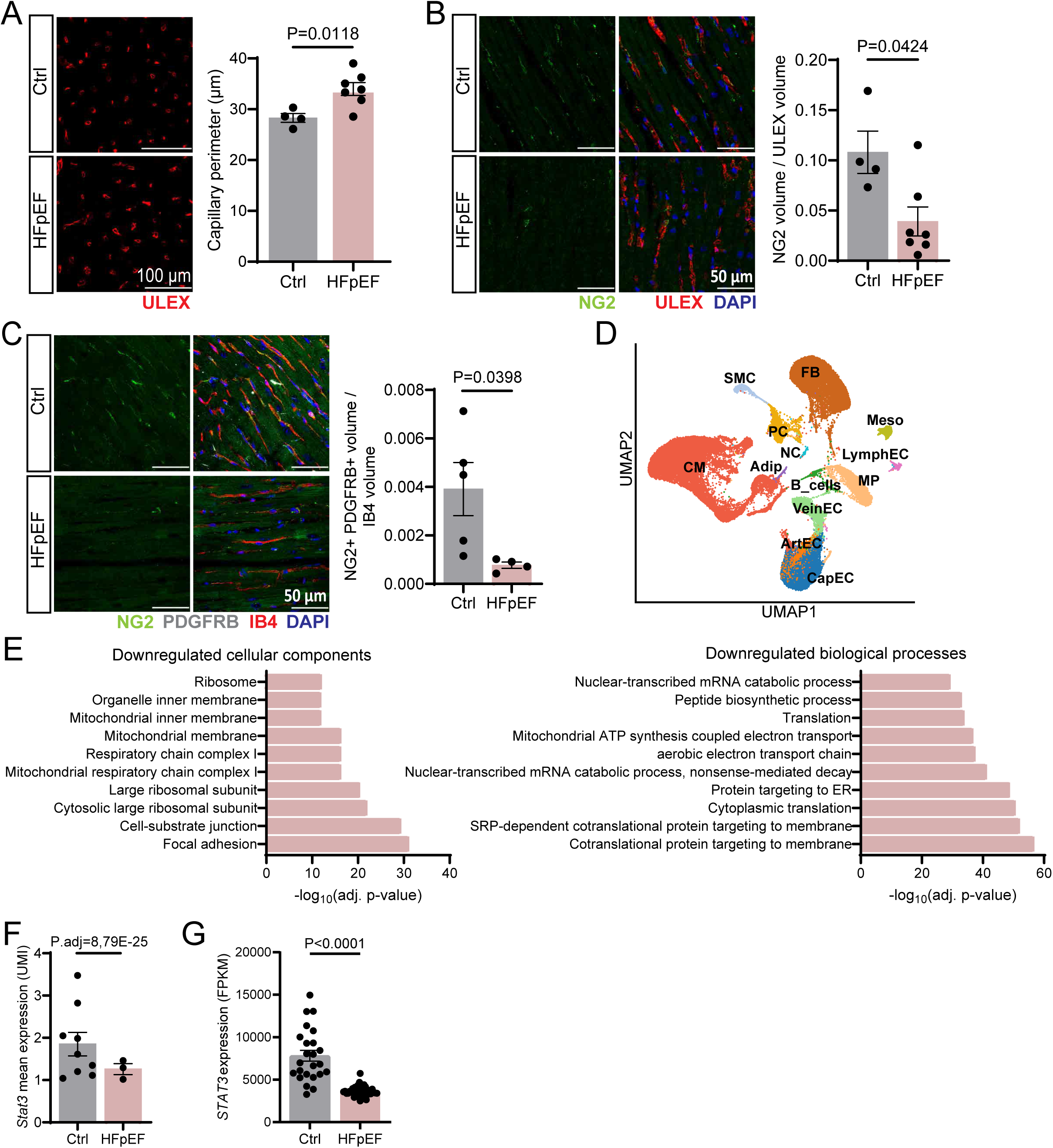
*STAT3* expression is reduced in HFpEF pericytes. (**A, B**) Immunofluorescence staining of control (Ctrl) and HFpEF patient biopsy. (**A**) Measurement of capillary perimeter. (**B**) Quantification of pericyte coverage normalized to the vasculature. N=4 for control and N=7 for HFpEF. Every data point represents one independent patient. Data are shown as mean ± S.E.M. P values were calculated using unpaired, two-tailed Student’s t-test. (**C**) Immunofluorescence staining of control and HFpEF mice. Quantification of pericyte coverage normalized to the vasculature. Pericytes were labelled with NG2 and PDGFRβ. N=5 for control and N=4 for HFpEF. Every data point represents one independent mouse. Data are shown as mean ± S.E.M. P value was calculated using unpaired, two-tailed Student’s t-test. (**D**) Uniform Manifold Approximation and Projection (UMAP) plot showing cell-type specific clustering of all data points from cardiac single-nuclei sequencing. we identified 13 individual cell types: Cardiomyocytes (CM), Artery (ArtEC), Vein (VeinEC), Capillary (CapEC) and Lymphatic (LymphEC) Endothelial Cells, B Cells, Macrophages (MP), Adipocytes (Adip), Fibroblasts (FB), Pericytes (PC), Smooth Muscle Cells (SMC), Meothelial cells (Meso), Neuronal cells (NC). (**E**) Gene Ontology (GO) enrichment analysis of significant differentially expressed genes in HFpEF pericytes. Represented are the top 10 downregulated cellular compartments and biological processes. (**F**) Scatter plot showing *Stat3* normalized gene expression values (unique molecular identifier, UMI) for the pericyte cluster in control and HFpEF pericytes. N=9 for control and N=3 for HFpEF. Every data point represents one independent mouse. Data are shown as mean ± S.E.M.P value was calculated using *bimod* test. (**G**) Scatter plot showing *Stat3* normalized gene expression values (fragments per kilobase of transcript per million mapped reads, FPKM) in control and HFpEF hearts. N=24 for control and N=41 for HFpEF. Every data point represents one independent patient. Data are shown as mean ± S.E.M. P value was calculated using Mann-Whitney test.

To gain insights into STAT3’s function in pericytes, we knocked down STAT3 in cultured human pericytes using siRNA (**Figure 2A**). First, we tested whether STAT3 depletion affected pericyte identity. Although *PDGFRB* and *CSPG4* expression remained unchanged (**Figure 2B**), we observed an increase in α smooth muscle actin (αSMA) (**Figure 2C**) and *COL3A1*, but not *COL1A1*, expression (**Figure 2D**) suggesting a shift towards a pro-fibrotic phenotype. Moreover, consistent with our HFpEF pericyte sequencing data, STAT3-deficient pericytes exhibited fewer focal adhesions per cell (**Figure 2E**), reduced *DES* expression (**Figure 2F**), and diminished cellular adhesion capacity (**Figure 2G**). Despite the effects on cellular adhesion and *DES* expression [23], *STAT3* knockdown did not significantly impact migration (**Supplementary Figure 2A-C**). Taken together, these observations suggest that *STAT3* deficiency impairs pericyte adhesion, potentially contributing to pericyte loss in HFpEF.

**Figure 2.**
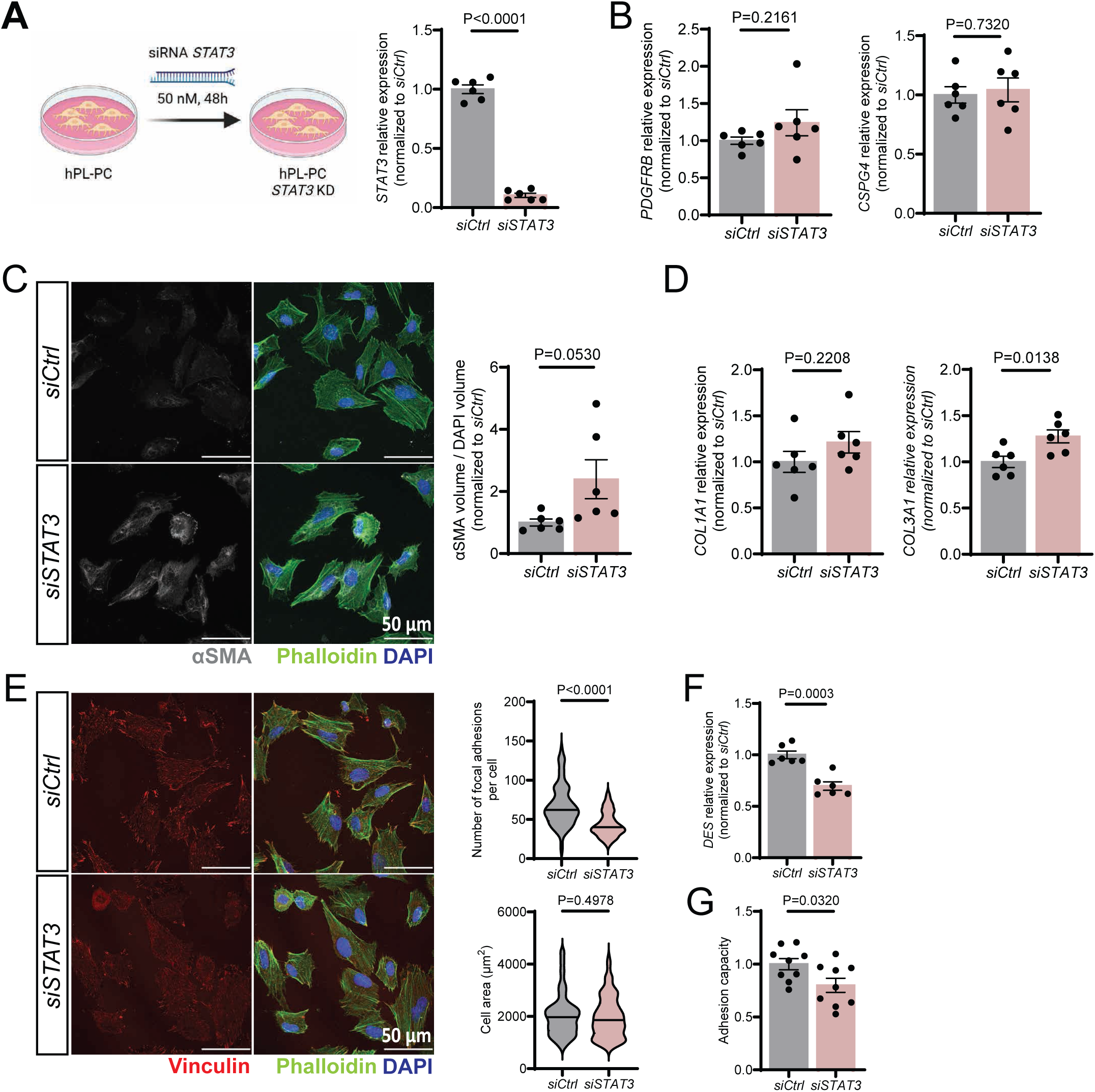
STAT3 deficiency compromises pericytes adhesion. (**A**) Scheme of the experimental design and RT-qPCR gene expression analysis of *STAT3* showing efficient *STAT3* silencing in pericytes. Every data point (n=6) represents an independent transfection. Data are shown as mean ± S.E.M. P values were calculated using unpaired, two-tailed Student’s t-test. (**B**) RT-qPCR gene expression analysis of *PDGFRB* and *CSPG4*. Every data point (n=6) represents an independent transfection. Data are shown as mean ± S.E.M. P values were calculated using unpaired, two-tailed Student’s t-test. (**C**) Immunofluorescence staining of control and STAT3 deficient pericytes. Every data point (n=6) represents an independent transfection. Data are shown as mean ± S.E.M. P values were calculated using unpaired, two-tailed Student’s t-test. (**D**) RT-qPCR gene expression analysis of *COL1A1* and *COL3A1*. Every data point (n=6) represents an independent transfection. Data are shown as mean ± S.E.M. P values were calculated using unpaired, two-tailed Student’s t-test. (**E**) Immunofluorescence staining of control and STAT3 deficient pericytes. Violin plots representing every analyzed cell in a total of N=6 independent transfections. P values were calculated using unpaired, Mann-Whitney test. (**F**) RT-qPCR gene expression analysis of *DES*. Every data point (n=6) represents an independent transfection. Data are shown as mean ± S.E.M. P values were calculated using unpaired, two-tailed Student’s t-test. (**G**) Adhesion capacity of control and STAT3 deficient pericytes. Every data point (n=9) represents an independent transfection. Data are shown as mean ± S.E.M. P values were calculated using unpaired, two-tailed Student’s t-test.

To determine whether STAT3 loss affects cellular proliferation, we performed a BrdU incorporation assay. Cell cycle analysis revealed no differences in the proportion of cells in G1 or S-phase but showed an accumulation in the G2/M phase (**Figure 3A**). Expression analysis of cell cycle regulators indicated an increase in the cell cycle inhibitor *CDKN1A*, but *TP53* levels were unaffected (**Figure 3B**). Furthermore, STAT3 deficient pericytes show increased dehydrogenase activity (**Figure 3C**) and telomere attrition (**Figure 3D**), suggesting that STAT3 loss may lead to a senescent phenotype [24–28]. β-Galactosidase staining further confirmed increased senescence in STAT3-deficient pericytes (**Figure 3E**).

**Figure 3.**
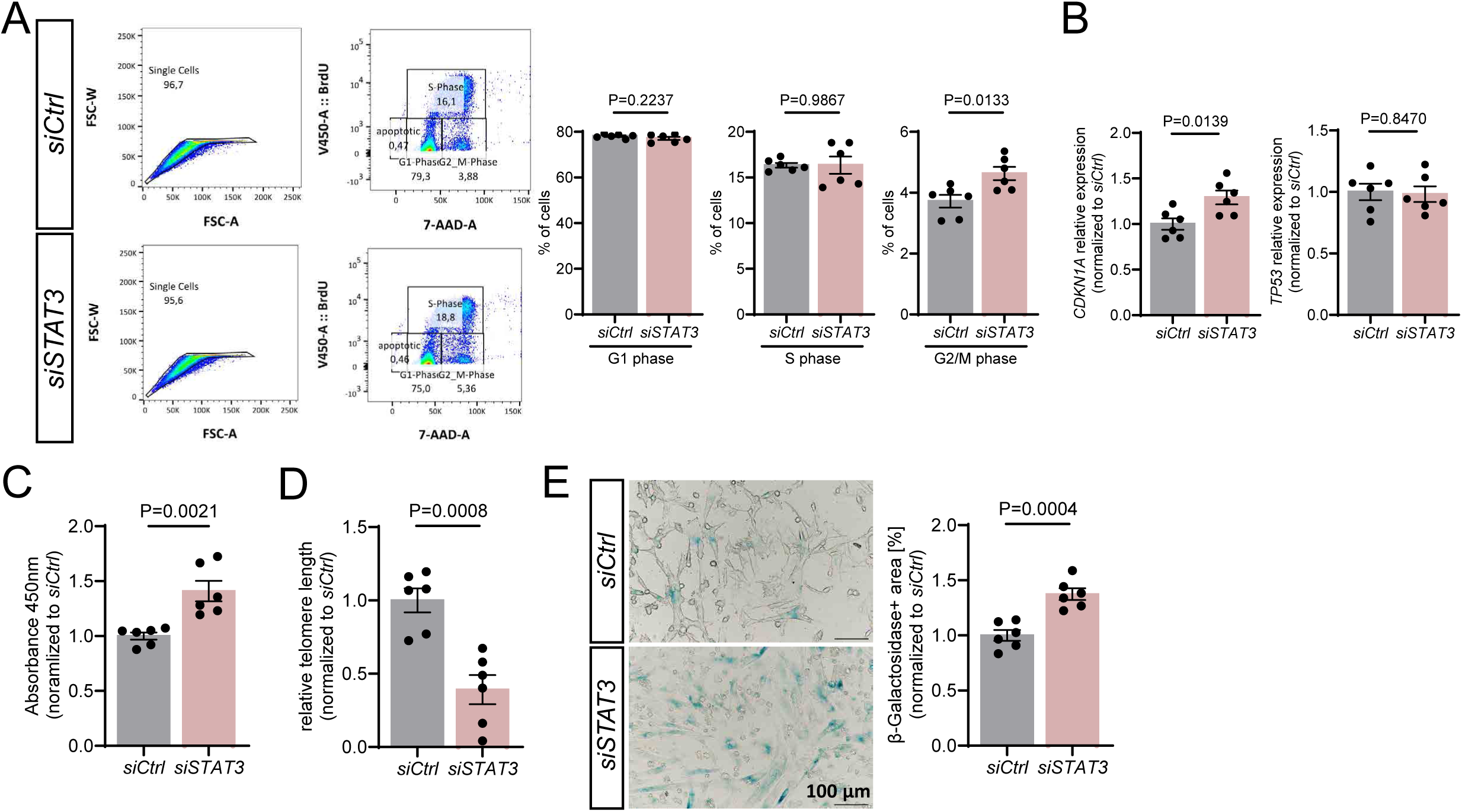
STAT3 deficiency induces cellular senescence in pericytes. (**A**) Representative FACS plots showing the gating strategy for BrdU proliferation assay. Cell cycle phases distribution differences in control and STAT3 deficient pericytes. (**B**) RT-qPCR gene expression analysis of *CDKN1A* and *TP53*. (**C**) Measurement of dehydrogenase activity in pericytes using CCK-8. (**D**) RT-qPCR analysis of relative telomere length in control and STAT3 deficient pericytes. (**E**) β-Galactosidase staining of control and STAT3 deprived pericytes and quantification of β-Galactosidase+ area (%). (**A**-**E**) Every data point (n=6) represents an independent transfection. Data are shown as mean ± S.E.M. P values were calculated using unpaired, two-tailed Student’s t-test.

To understand the molecular mechanism by which STAT3 deficiency in pericytes compromises cellular adhesion and induces senescence, we performed bulk-RNA-sequencing on STAT3-deficient pericytes. Gene expression analysis revealed 1563 significant DEGs (P<0.05; logFoldChange>0.025), including 860 upregulateda and 703 downregulated genes. GO analysis of regulated pathways showed dysregulation in genes associated with senescence (**Figure 4A**), confirming our previous observations. In particular, STAT3-deficient pericytes showed increased expression of cell cycle inhibitors such as *CDKN1A*, *CDKN1B* or *CDKN2B* (**Figure 4B**). Consistent with the transcriptomic signature of pericytes in HFpEF mouse hearts, GO analysis of cellular components revealed dysregulation in genes related to focal adhesion and cell-substrate junctions (**Figure 4C**), including focal adhesion-associated genes like *ARPC5L* [29], substrate adhesion genes such as *RAB2* [30], and cell-to-cell junction genes like *GJA1* **(Figure 4D)**.

**Figure 4.**
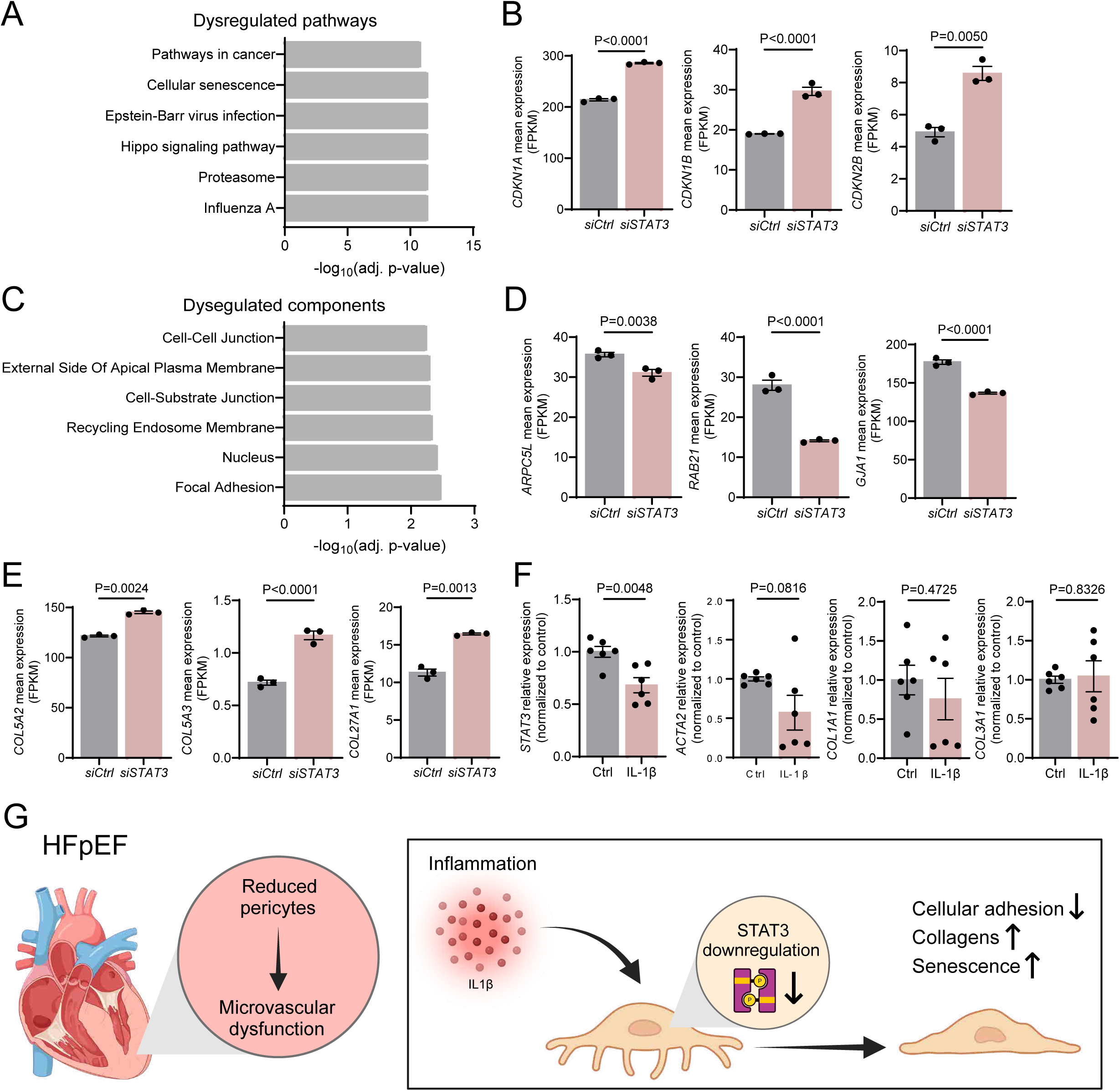
STAT3 knockdown induces a transcriptional signature similar to HFpEF. (**A**) Gene Ontology (GO) enrichment analysis of significant differentially expressed genes in STAT3 deficient pericytes. Represented are the top 6 dysregulated pathways. (**B**) Scatter plot showing *CDKN1A*, *CDKN1B* and *CDKN2B* normalized gene expression values (fragments per kilobase of transcript per million mapped reads, FPKM) in control and STAT3 deficient pericytes hearts. Every data point (n=3) represents an independent transfection. Data are shown as mean ± S.E.M. P values were calculated using cuffdiff test. (**C**) Gene Ontology (GO) enrichment analysis of significant differentially expressed genes in STAT3 deficient pericytes. Represented are the top 6 dysregulated cellular components. (**D, E**) Scatter plot showing *ARPC5L*, *RAB21*, *GJA1, COL5A2, COL5A3* and *COL27A1* normalized gene expression values (fragments per kilobase of transcript per million mapped reads, FPKM) in control and STAT3 deficient pericytes hearts. Every data point (n=3) represents an independent transfection. Data are shown as mean ± S.E.M. P values were calculated cuffdiff test. (**F**) RT-qPCR gene expression analysis of *STAT3, ACTA2, COL1A1* and *COL3A1* in pericytes upon IL-1β treatment. Every data point (n=6) represents an independent transfection. Data are shown as mean ± S.E.M. P values were calculated using unpaired, two-tailed Student’s t-test. (**G**) Graphical summary of the study

Increased expression of extracellular matrix collagens, including *COL5A2*, *COL5A3*, and *COL27A1* (**Figure 4E**), further confirmed the pro-fibrotic activation of pericytes upon STAT3 deletion. These findings suggest that STAT3 represses a senescent gene programme and maintains focal adhesion signalling in pericytes.

Finally, we explored the cause of reduced STAT3 expression in HFpEF pericytes. As inflammation is a major hallmark and driver of HFpEF [31,32], and IL-1β is a key cytokine implicated in HFpEF development [33], we investigated whether IL-1β modulates STAT3 expression in pericytes. IL-1β treatment for 24 hours significantly reduced *STAT3* expression in pericytes (**Figure 4F**) suggesting inflammation as a driver of *STAT3* downregulation in pericytes. Interestingly, IL-1β-treated pericytes did not exhibit a pro-fibrotic phenotype, as expression levels of *ACTA2*, *COL1A1* and *COL3A1* were not significantly altered (**Figure 4F**), indicating that pro-fibrotic activation may occur downstream of STAT3 downregulation.

## Discussion

In the present study, we have studied the molecular alterations in pericytes in the context of HFpEF. We identified reduced STAT3 expression in pericytes as a characteristic of HFpEF in both human and a murine model of the disease. Furthermore, we demonstrated that STAT3 deletion in pericytes recapitulates several HFpEF-related features, such as decreased focal adhesion and cellular adhesion loss. In line with our findings, STAT3 localises to focal adhesions in cancer cells [34] and, upon activation, contributes to in tumour metastasis through increased expression of cell adhesion molecules [35]. Additionally, STAT3 activity modulates the DNA damage response [36], and cells lacking STAT3 are more sensitive to oxidative stress [37]. DNA damage in pericytes has already been observed in other pathological disorders characterized by pericyte coverage reduction. In a model of diabetic retinopathy, enhanced expression of thioredoxin-interacting protein (TXNIP) induces oxidative stress, DNA damage, and pericyte loss [38]. In neuronal ceroid lipofuscinosis, oxidative DNA damage reduces the viability of brain pericytes inducing microvascular dysfunction impairing blood-brain barrier [39]. Reduced pericyte coverage in the brain is one of the structural changes that contributes to blood-brain barrier disruption during HIV infection. Interestingly, human brain vascular pericytes infected with HIV-1 show significant accumulation of DNA damage [40].

Based on this, we propose that inflammation in HFpEF may reduce the expression of STAT3 in pericytes, causing DNA damage, telomere shortening, increased dehydrogenases activity and cellular senescence. This may be accompanied by focal adhesion disassembly, resulting in decreased pericyte coverage of myocardial capillaries (**Figure 4G**). In conclusion, our study identifies pericyte STAT3 reduction as a marker of HFpEF and highlights the importance of pericytes in maintaining cardiac homeostasis. Our findings provide the first evidence that STAT3 is essential for sustaining pericyte function, preventing pericyte loss, pro-fibrotic gene expression, and cellular senescence.

## Acknowledgments

L.R.V. would like to thank Anita Tamiato for her technical and mental support.

## Sources of funding

This study was supported by the DFG SFB1531 project number 456687919. Project B5 to GL and the BMBF-funded project DZHK RheinMain “Cellular Heterogeneity” and THERANOVA (supported by the State of Hesse) to SD.

## Autor contributions

L.R.V. performed in vitro and in vivo experiments and data analysis. A.F. produced the murine HFpEF histological slides. T.P. and J.W. performed bulk RNA sequencing. M.R.J. and D.J. performed bioinformatic analysis. M.M. analysed FACS data. B.K. Provided the mouse HFpEF sequencing data. V.P, F.E. and E.N. provided the human HFpEF biopsies. S.D. Obtained funding and provided supervision. G.L. conceived and designed the experiments, wrote the manuscript, obtained funding for the project and supervised the project.

## Disclosures

None.

---

**Supplementary Figure 1.**
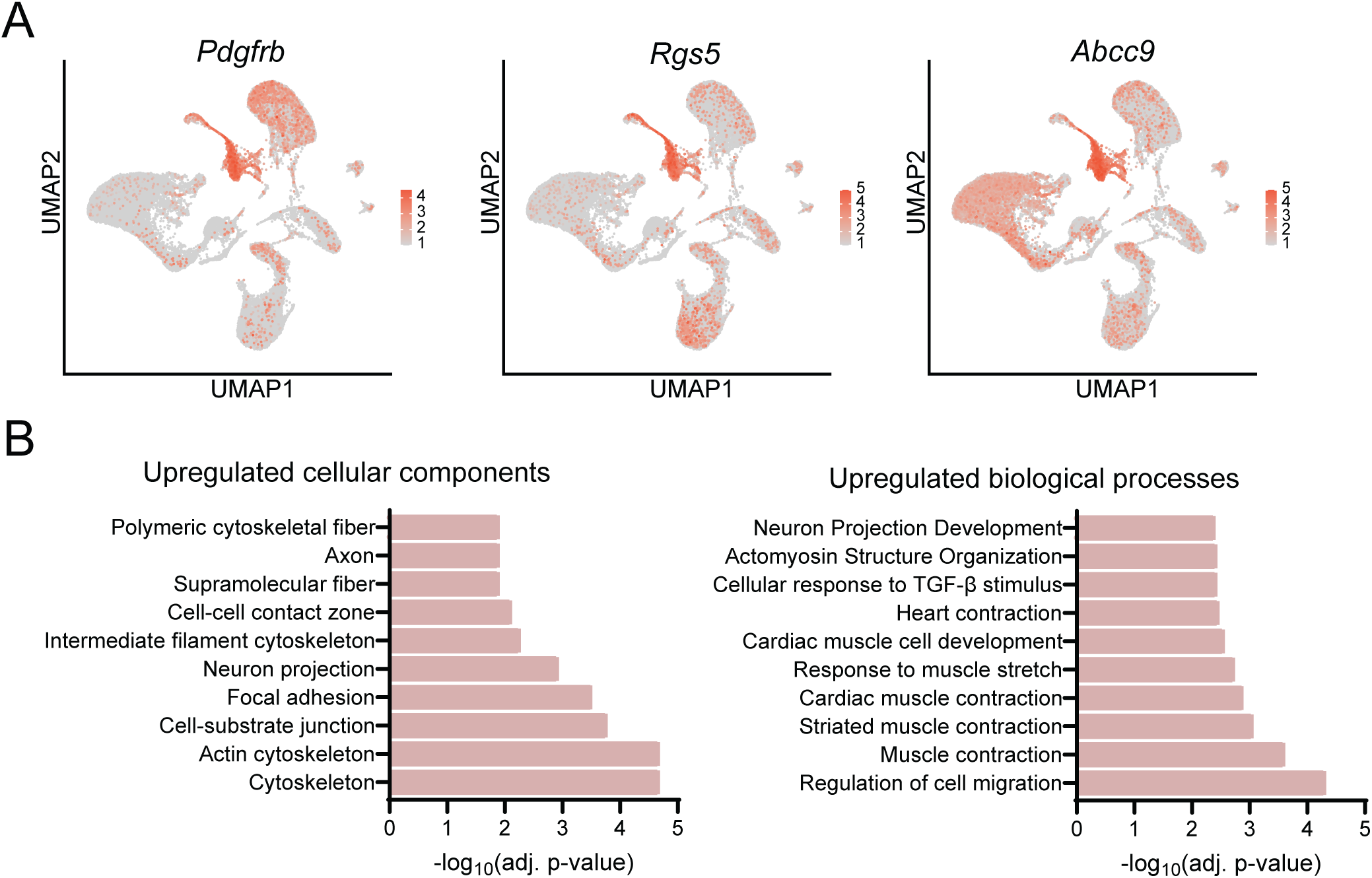
Gene Ontology (GO) enrichment analysis of significant differentially expressed genes. (**A**) Feature plot showing gene expression of pericyte marker genes (*Pdgfrb*, *Rgs5*, *Abcc9*). The coloured scale bar indicates the log-normalized gene expression level. (**B**) Gene Ontology (GO) enrichment analysis of significant differentially expressed genes in HFpEF pericytes. Represented are the top 10 upregulated cellular compartments and biological processes.

**Supplementary Figure 2.**
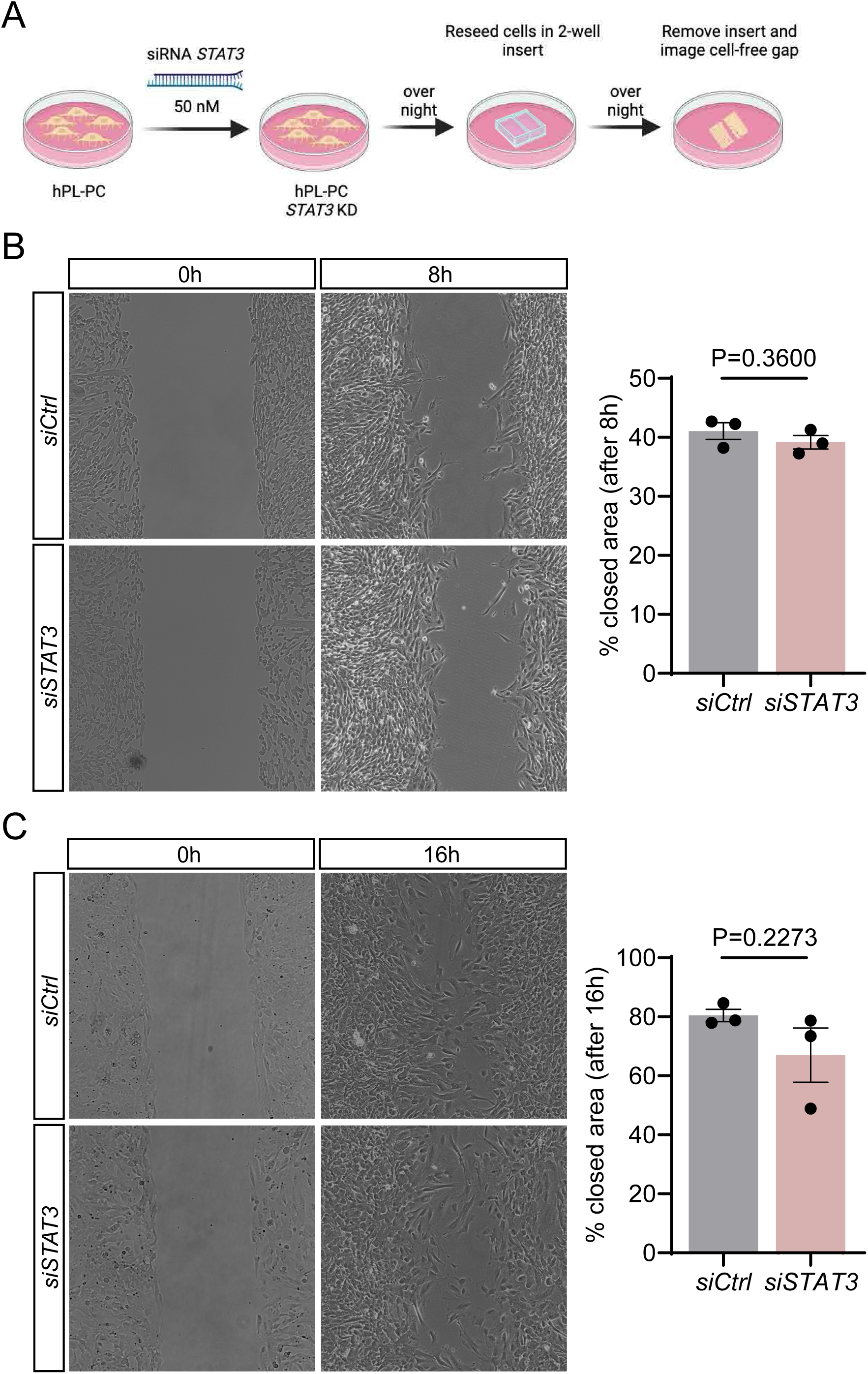
STAT3 deficiency does not significantly affect cellular migration. (**A**) Scheme of experimental design. (**B**, **C**) Quantification of migrated area by control and STAT3 deficient pericytes after 8h or 16h. Every data point (n=3) represents an independent transfection. Data are shown as mean ± S.E.M. P values were calculated using unpaired, two-tailed Student’s t-test.

## Notes

### Competing Interest Statement

The authors have declared no competing interest.

